# “The anti-angiogenic compound axitinib demonstrates low toxicity and anti-tumoral effects against medulloblastoma”

**DOI:** 10.1101/2020.09.18.301028

**Authors:** Marina Pagnuzzi-Boncompagni, Vincent Picco, Valérie Vial, Victor Planas-Bielsa, Ashaina Vandenberghe, Renaud Grépin, Jérôme Durivault, Christopher Montemagno, Sonia Martial, Jérôme Doyen, Julie Gavard, Gilles Pagès

**Affiliations:** Centre Scientifique de Monaco, Biomedical Department, 8 Quai Antoine Ier, MC-98000 Monaco, Principality of Monaco; Centre Scientifique de Monaco, Polar Biology Department, 8 Quai Antoine Ier, MC-98000 Monaco, Principality of Monaco; University Nice Cote d’Azur, Institute for Research on Cancer and Aging of Nice (IRCAN), CNRS UMR 7284; INSERM U1081, Centre Antoine Lacassagne, France; Department of Radiation Oncology, Centre Antoine-Lacassagne, University of Côte d’Azur, Fédération Claude Lalanne, Nice, France; Team SOAP, CRCINA, INSERM, CNRS, Université de Nantes, Nantes, France; Integrated Center of Oncology, St-Herblain, France

## Abstract

Evolution of medulloblastoma (MB) treatments has increased the 5-year overall survival of to more than 70%. However, an increasing number of survivors face severe long-term adverse effects and associated morbidity due to multimodal treatments particularly harsh for the younger patients. Chemotherapeutic compounds inducing less adverse effects are key to improving the care of MB patients. The preclinical relevance of last generation anti-angiogenic compounds deserves to be fully assessed. Among these, axitinib showed the highest selectivity index for MB cells, efficiently reduced the growth rate of experimental tumors and led to less toxicity towards normal cells than did a reference treatment. In vivo, axitinib did not lead to acute toxicity in very young rats and was able to cross the blood brain barrier. Analysis of public databases shows that high expression of axitinib targets are of poor prognosis. Altogether, our results suggest that axitinib is a compelling candidate for MB treatment.

## INTRODUCTION

Cancer is the second cause of mortality during childhood in high income countries, behind accidental death. Though high during the last decades, the decrease rate of child cancer mortality tends to reach a plateau ^1^. This suggests that the use of current anti-cancer drugs is reaching maximum optimization. New improvements in childhood cancer care will therefore need the development of new therapeutic approaches.

Cancers of the central nervous system are the second most prevalent childhood tumors after hematologic cancer ^2^. MB is the most prevalent brain cancer in children and infants and accounts for 15-20% of childhood nervous system tumors ^3^. The current therapeutic approach comprises surgical removal of the tumor, cranio-spinal radiation therapy (RT) and chemotherapy. It is based on the subdivision of patients into standard or high risk groups based on the presence of metastases, age, extent of postsurgical residual disease and histology [2, 29, 30]. Patients belonging to the standard risk group receive lower doses of radio- and chemotherapy in order to limit the deleterious effects of the treatments as much as possible.

Although optimization of MB treatment has led to an increase in the long-term survival and a decrease of recurrence, the rate of deleterious late effects (occurring more than 5 years after diagnosis) also significantly increased from the 1970s to the 1990s ^4^. The severe late outcomes include occurrence of a secondary neoplasm, severe psychological disorders and cardiac toxicity ^4-6^. These are suggested to be due to the introduction of adjuvant systemic chemotherapies in the treatment of MB during the 1980s ^4^. A challenge in modern MB therapy therefore lies in finding new treatments that allow maintenance or increase of survival while reducing adverse effects.

Recent advances in the genetic characterization of the disease have led to the classification of MBs into subtypes: the wingless (Wnt), the sonic hedgehog (Shh), and the more similar though molecularly distinguishable groups 3 and 4 [2]. MB subgrouping could thus be used to orient the therapeutic approach. Indeed, targeted therapies may present less off-target effects than classical genotoxic chemotherapies and therefore represent a promising approach to increase tolerability of the treatments and reduce deleterious side effects ^7^. Among these therapies, antagonists targeting Smoothened, the receptor of Hh ligands, such as Sonidegib (LDE225) and Vismodegib (GDC-0449) were considered extremely promising for the treatment of Shh subgroup MBs. Unfortunately, the first interventional studies using these drugs on pediatric Shh subgroup MB patients showed limited response rates associated with permanent defects in bone growth [31-33].

Interestingly, increased angiogenesis is associated with the most aggressive MBs (group 3) ^8^. Moreover, direct cytotoxic effects towards the tumor cells aberrantly expressing the targets of anti-angiogenic drugs have been observed ^9^. This suggests that anti-angiogenic treatments might be of interest for MBs. Few studies have been conducted with anti-angiogenic targeted compounds in a pediatric setup. The VEGF-targeting monoclonal antibody bevacizumab was used in combination with a variety of non-targeted chemotherapeutic agents for the treatment of pediatric patients with solid tumors, including some brain tumors ^10-14^. Despite a good tolerability of bevacizumab, no modification of event-free, progression-free or overall survival was observed. Previous work from our lab has demonstrated that the multi-target anti-angiogenic tyrosine kinase inhibitors (TKi) are directly toxic for cancer cells in addition to their in vivo anti-angiogenic activity [9, 34-36]. Amongst these, the TKi sunitinib has been tested in pediatric setups. This study concluded a lack of benefit and, more importantly, an increase in the occurrence of adverse events with the sunitinib treatment ^15^. The efficacy of axitinib, another multi-target anti-angiogenic TKi, to treat children with solid tumors has also recently been evaluated [16]. A phase I study determined the maximum tolerated and recommended dose in children presenting refractory or recurrent tumors (2.4mg/m^2^) [16]. Finally, cabozantinib is currently under clinical investigations to treat pediatric cancers but no results are available to our knowledge ^16, 17^.

Our study aimed at determining the efficacy of anti-angiogenic treatments in MB preclinical models. We found that sunitinib, cabozantinib and axitinib effectively kill MB cells in vitro. However, axitinib presented the best selectivity towards cancer cells when compared to primary normal cells. Axitinib also displayed in vivo efficacy. Finally, in an in vivo model of regulatory toxicity, no acute toxicity of this compound towards young and growing rats was observed ^18^. Moreover, axitinib was detected in the brain of the animals and was able to permeabilize an in vitro blood-brain barrier (BBB) model, which strongly suggests that it could efficiently reach brain tumors. Although our results should be considered with caution, they suggest that axitinib represents an option for patients in therapeutic impasse.

## RESULTS

### Among the available anti-angiogenic compounds, axitinib is the most selective for MB cells

In addition to their effects on tumor angiogenesis, anti-angiogenic compounds often present cytotoxicity towards tumor cells ^9^. In order to test if this was the case with MB cells, we compared the effect of three anti-angiogenic compounds (axitinib, cabozantinib and sunitinib) with that of a chemotherapy used in clinics to treat high risk MBs ^19^ (the Carboplatin/Etoposide combination, hereafter referred to as the “reference treatment”). In parallel, the toxicity of the compounds towards non-tumor cells was evaluated on primary astrocytes (C8-D1A) and fibroblasts (HDF). First, we determined the EC50 of each compound on these cell lines (Table 1). We also expressed the degree of selectivity of the compounds by calculating a selectivity index (SI) (Table 2) ^20^. SI was determined as the ratio of the EC50 of pure compound in a normal cell line and the EC50 of the same pure compound in MB cell lines. We showed that sunitinib, axitinib and cabozantinib impact Shh group (DAOY), group 3 (D458 and HD-MB03) and group 4 (CHLA-01-MED) MB cell lines ^21^ in a way comparable to the reference treatment (Table 1). However, the EC50 of axitinib on astrocytes and fibroblasts was extremely high, resulting in a much higher SI as compared to the other treatments, including the reference treatment (Table 2). Second, we evaluated the effect of each compound on the cell viability and proliferation of all five MB cell lines ^21^ and of the non-tumoral primary cells used in the previous experiments (Fig. 1). Cells were exposed to the compounds for 48 hours and their viability was measured by a propidium iodide incorporation technique (Fig. 1a). Axitinib, cabozantinib and sunitinib significantly decreased the viability and proliferation of most MB cells (Fig. 1a-c). Furthermore, axitinib, cabozantinib and the reference treatment had no effect on the viability of the HDF and C8-D1A non-tumor primary cells while sunitinib strongly induced the death of C8-D1A astrocytes (Fig. 1a and b). In order to get insight into the dynamics of the effect of each compound, cell proliferation was evaluated on cells continuously treated during four days (Fig. 1c). Cabozantinib and sunitinib decreased DAOY, HD-MB03 and D283 cell proliferation approximately by half, whereas axitinib and the reference treatment completely inhibited cell proliferation. Conversely, axitinib only had a moderate impact on HDF proliferation while it strongly prevented the proliferation of C8-D1A cells (Fig. 1c). These results suggest that the anti-angiogenic compounds we tested display a direct effect on MB cells. However, Axitinib was the only anti-angiogenic compound that displayed a selective effect towards MB versus non-tumor cells in MTT assays. Moreover, our results show that axitinib has a selective inhibitory effect on tumor versus non-tumor cells.

**Table 1:** Toxicity of antiangiogenic compounds and chemotherapeutic agents towards non-tumoral and MB cells. The EC50 (µM) values for various compounds determined by MTT test after 48 hours exposure of non-tumoral and MB cell lines are presented (values are mean IC50 +/- standard error to the mean from at least three independent experiments).

|  | Non-tumoral cells |  | Medulloblastoma cells |  |  |  |
| --- | --- | --- | --- | --- | --- | --- |
| EC50 (μM) | HDF | C8-D1A | D458 | DAOY | HD-MB03 | CHLA-01-Med |
| **Sunitinib** | 1.8+/-0.04 | 5.7+/-2.2 | 4.2+/-0.3 | 7.9+/-5.5 | 4.6+/-0.3 | 5.3 |
| **Cabozantinib** | 4.1+/-1.9 | 4+/-2.2 | 5.1+/-1.6 | 8.3+/-1.2 | 0.78+/-0.2 | 26.5 |
| **Carbo/Eto** | 1.6+/-1.1 | 3+/-0.8 | 0.56+/-0.7 | 1+/-0.4 | 0.16+/-0.07 | n.d. |
| **Axitinib** | **>100** | **>100** | 0.49+/-0.4 | 2.3+/-1.5 | 0.47+/-0.2 | 14.6 |

**Table 2:** Specificity indexes for three antiangiogenic compounds and reference chemotherapy in normal and MB cells. Specificity indexes were determined by calculating the ratio between the IC50 of a given drug for either human dermal fibroblasts (HDF) or murine primary astrocytes (C8-D1A) and the IC50s of the same drug for each of the MB cell lines (D458, DAOY, HDMB and CHAL-01-Med).

| Specificity index | HDF |  |  |  | C8D1A |  |  |  |
| --- | --- | --- | --- | --- | --- | --- | --- | --- |
|  | D458 | DAOY | HDMB | CHLA-01-Med | D458 | DAOY | HDMB | CHLA-01-Med |
| **Sunitinib** | 0.43 | 0.23 | 0.39 | 0.34 | 1.36 | 0.72 | 1.24 | 1.08 |
| **Cabozantinib** | 0.80 | 0.49 | 5.26 | 0.15 | 0.78 | 0.48 | 5.13 | 0.15 |
| **Carbo/Eto** | 2.86 | 1.60 | 10.00 | n.d. | 5.36 | 3.00 | 18.75 | n.d. |
| **Axitinib** | >200 | >40 | >200 | >6 | >200 | >40 | >200 | >6 |

**Figure 1:**
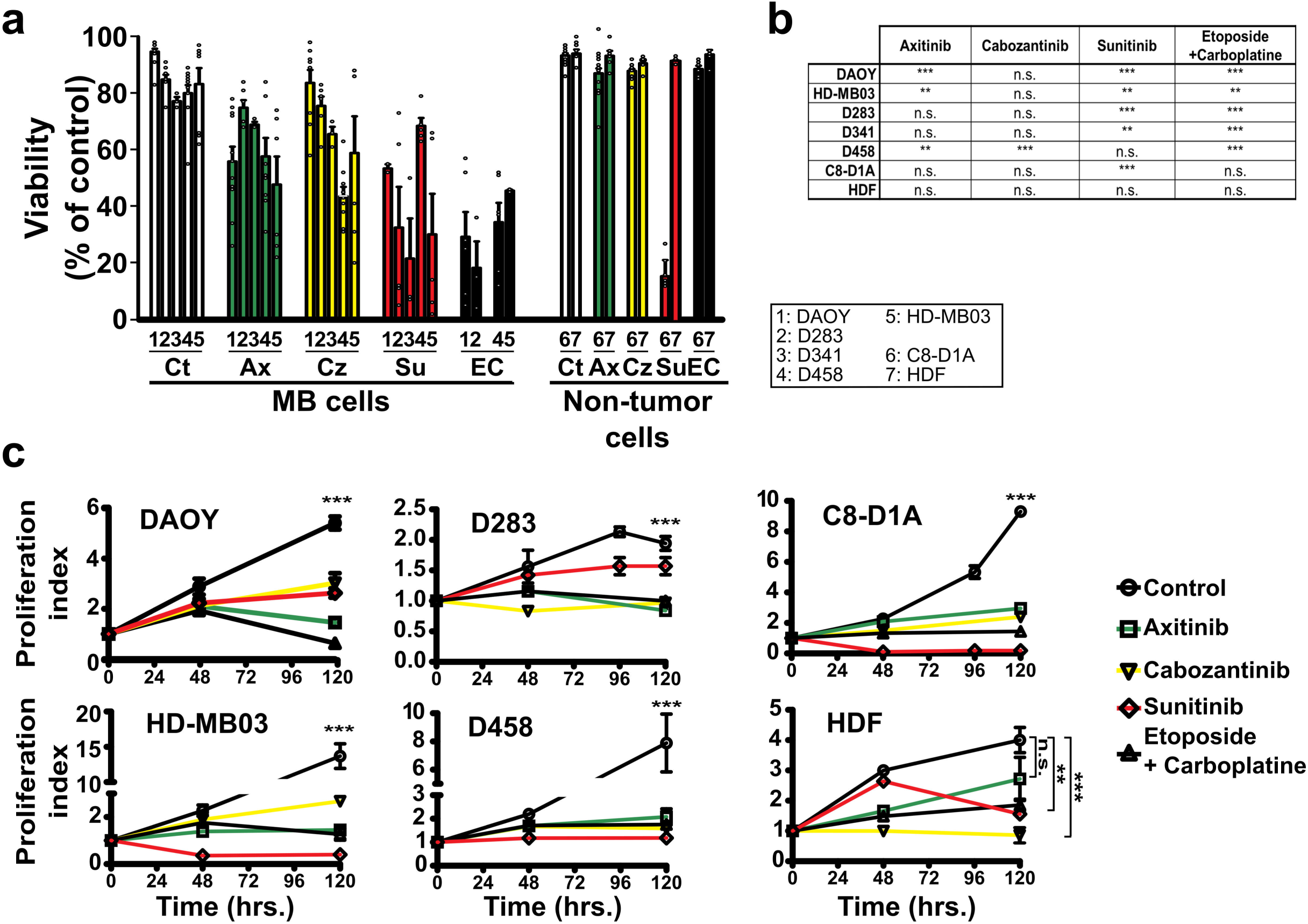
Antiangiogenic compounds strongly impair the viability and proliferation of MB cells. (a) Viability of the indicated cell lines treated for 48 hours with axitinib (Ax, green), cabozantinib (Cz, yellow), sunitinib (Su, red) or a combination of etoposide and carboplatin (EC, black). All the compounds were dissolved in DMSO, the amount of which was adjusted to be the same in every condition. Control conditions (Ct, white) are also treated with the same amount of DMSO. Histograms represent the mean +/- SEM; each independent data point is represented (white circles). (b) Table showing the statistical significances of the viability results for each compound compared to the control condition (***: p<0.001, **: p<0.01, n.s.: non-significant, Student’s t-test comparing at least three independent experiments). (c) Proliferation index of MB and normal cell lines continuously treated with the indicated concentrations of each compound (***: p<0.001, **: p<0.01, n.s.: non-significant, one-way Anova test comparing controls to the other experimental conditions).

### Axitinib leads to low toxicity towards non-tumoral brain cells

Considering our results and previous studies showing a potential efficiency of axitinib on MB cells ^22^ and no efficiency of sunitinib on pediatric brain tumors ^15^ we chose to focus our study on axitinib. To further characterize the differential effect of axitinib and etoposide/carboplatin reference treatment on MB and normal cells, we stained HD-MB03, DAOY and C8-D1A cells with different fluorescent probes. MB cells and C8-D1A astrocytes were co-cultured and treated with axitinib or etoposide/carboplatin. Both treatments resulted in a 40 to 60% enrichment of the proportion of astrocytes relative to the total number of cells as quantified by FACS (Fig. 2a and b). The long-term proliferation in a cumulative population doubling (C.P.D.) experiment (Fig. 2c) showed that continuous treatment of DAOY and HD-MB03 cells with both compounds strongly impaired their proliferation (Fig. 2c). Continuous treatment of non-tumor cells C8-D1A and HDF also impaired their proliferation, although axitinib had a milder effect on C8-D1A cells compared to the reference treatment (Fig. 2c). We performed the same experiment with a transient 48 hour treatment followed by removal of the compounds (Fig. 2c). Axitinib and etoposide/carboplatin were sufficient to reduce proliferation of MB cells by more than 80%. Proliferation of normal cells was also impaired in these conditions, although to a significantly lower extent with axitinib as compared to etoposide/carboplatin (Fig. 2c). Thus, this data confirms that axitinib selectively impairs tumor cell survival and further suggests that axitinib leads to lower toxicity towards non tumoral brain cells than the etoposide/carboplatin chemotherapy reference treatment.

**Figure 2:**
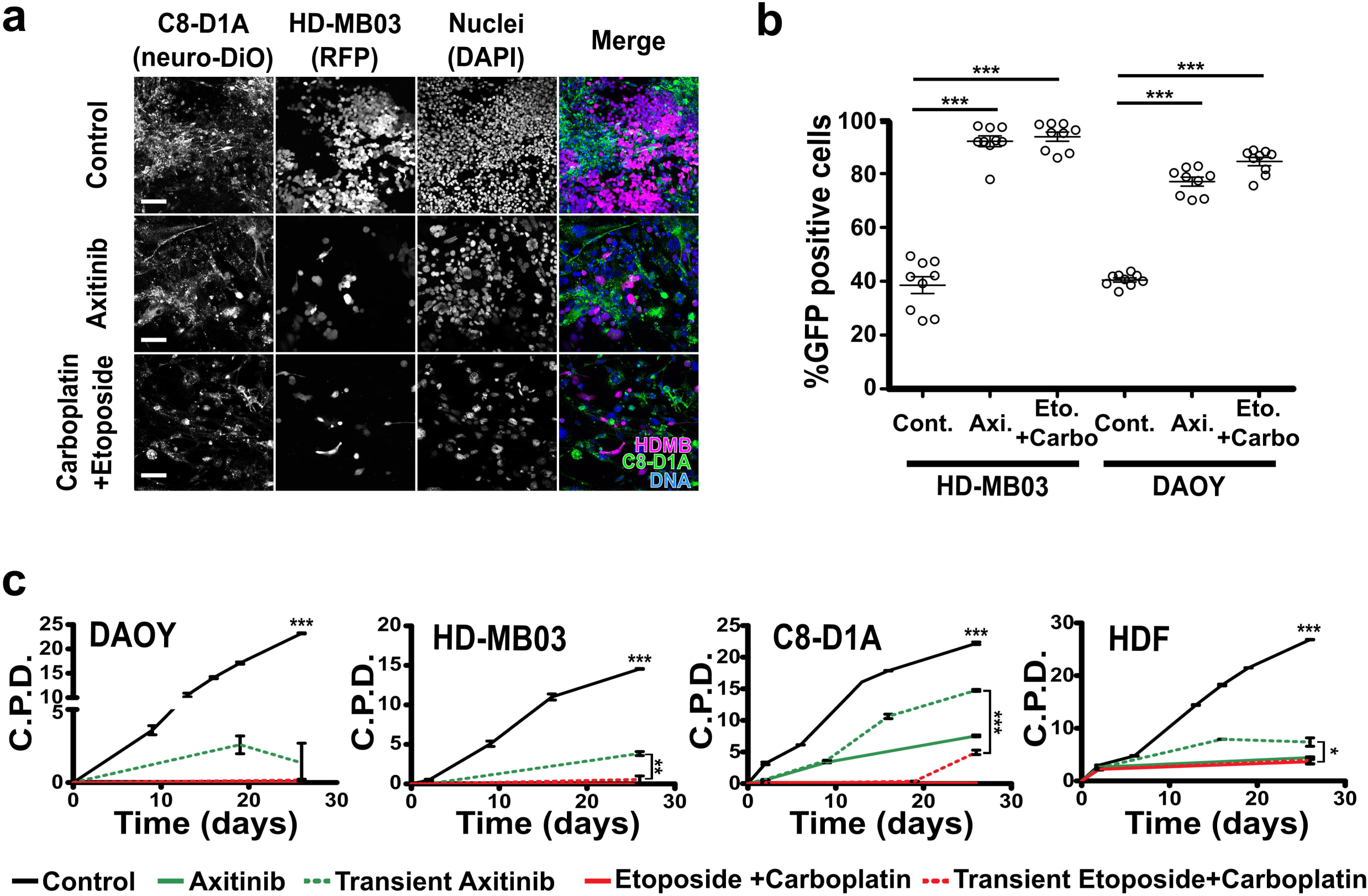
Axitinib selectively impacts proliferation of MB cells in 2D models. (a) Cumulative population doublings (C.P.D.) of the indicated MB and normal cell lines continuously treated with 5µM axitinib (continuous green lines), a combination of 1µM etoposide and 1.6µM carboplatin (EC, continuous red lines) or transiently treated for 48 hours with the same compound concentrations (dotted lines) (data points represent the mean +/- SEM of a representative experiment; ***: p<0.001, **: p<0.01, *: p<0.05, one-way Anova test, results are statistically non-significant unless otherwise stated). (b) Confocal images of RFP-expressing MB cell (HDMB, magenta) and neuro-DiO-stained primary astrocyte (C8-D1A, green) cocultures transiently treated for 4 days with 5µM Axitinib or a combination of 1µM etoposide and 1.6µM carboplatin (Hoechst 33342 nuclear DNA counterstain is showed in blue; scale bars: 100µm). (c) FACS quantification of neuro-DiO positive cells relative to the total number of cells (horizontal lines and errors bars represent the mean +/- SEM and each individual data point of three independent experiments are plotted; ***: p<0.001, one-way Anova test, results are statistically non-significant unless otherwise stated).

### Axitinib reduces MB cell proliferation in 3D cultures

We next generated spheroids combining fluorescent MB and non-tumor cells to test the selectivity of axitinib on structures mimicking experimental tumors. Axitinib and etoposide/carboplatin treatments resulted in depletion of most of the tumor cells while the normal cells were preserved (Fig. 3a). The growth over five days of spheroids formed with four different MB cell lines originally isolated form Shh (DAOY), group 3 (HD-MB03, D283, CHLA-01-MED) and group 4 (D458) subgroups of MB was evaluated (Fig.3b and Supplementary Fig. S1). Axitinib abolished 3D growth of MB cells in a way comparable to the reference treatment. Interestingly, a combination of half-dose axitinib/etoposide (2.5µM and 0.5µM respectively) in the DAOY and HD-MB03 spheroids (Fig.3b) reduced spheroid growth more efficiently than the etoposide/carboplatin treatment. These results favor the use of axitinib in combination with the chemotherapy reference compound etoposide.

**Figure 3:**
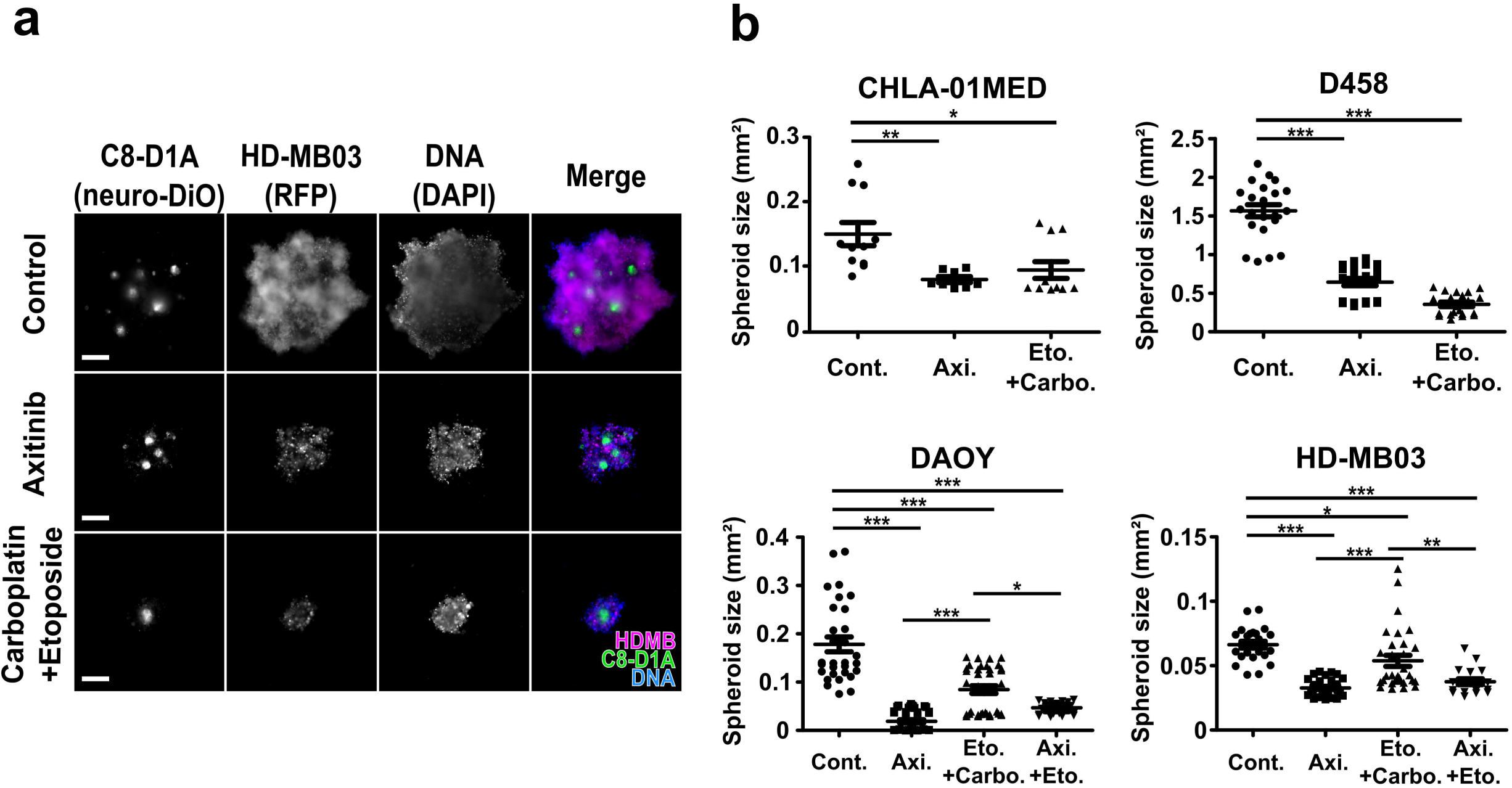
Axitinib selectively impacts proliferation of MB cells in 3D models. (a) Mixed spheroids generated with neuro-DiO-stained astrocytes (C8-D1A, green) and RFP-expressing MB cells (HD-MB03, magenta) treated with 5µM axitinib (Ax.) or a combination of 1µM etoposide and 1.6µM carboplatin (EC) (scale bars: 250µm). (b) Dot plots showing the endpoint size measurements of spheroids generated with the indicated MB cell lines and continuously treated with 5µM axitinib (green lines) or a combination of 1µM etoposide and 1.6µM carboplatin (red lines). Controls were all treated with a concentration of DMSO corresponding to the one used as vehicle for the drugs (a representative experiment of at least 3 independent experiments is presented; data points represent the mean +/- SEM; ***: p<0.001 one-way Anova test, results are non-statistically significant unless otherwise stated).

### Axitinib decreases the viability of radiation-resistant MB cells

Resistance to the treatment is one of the major causes for the loss of control of diseases and aggressiveness of relapsed tumors. We tested the effect of axitinib on DAOY and HD-MB03 cells rendered resistant to radiotherapy after multiple exposures of low-dose radiations (courtesy of Dr. Sonia Martial, unpublished). Treatment with axitinib, etoposide/carboplatin and axitinib/etoposide for 48 hours reduced the viability of control and radiation-resistant cells to a comparable extent (Fig.4a). More strikingly, all the treatments efficiently abolished the proliferation of control and radiation-resistant cells (Fig.4b). This shows that the acquisition of radio-resistance by MB cells does not confer resistance of these cells to either the reference treatment or axitinib. This result suggests that tumors progressing under radio-therapy may be efficiently treated with axitinib.

**Figure 4:**
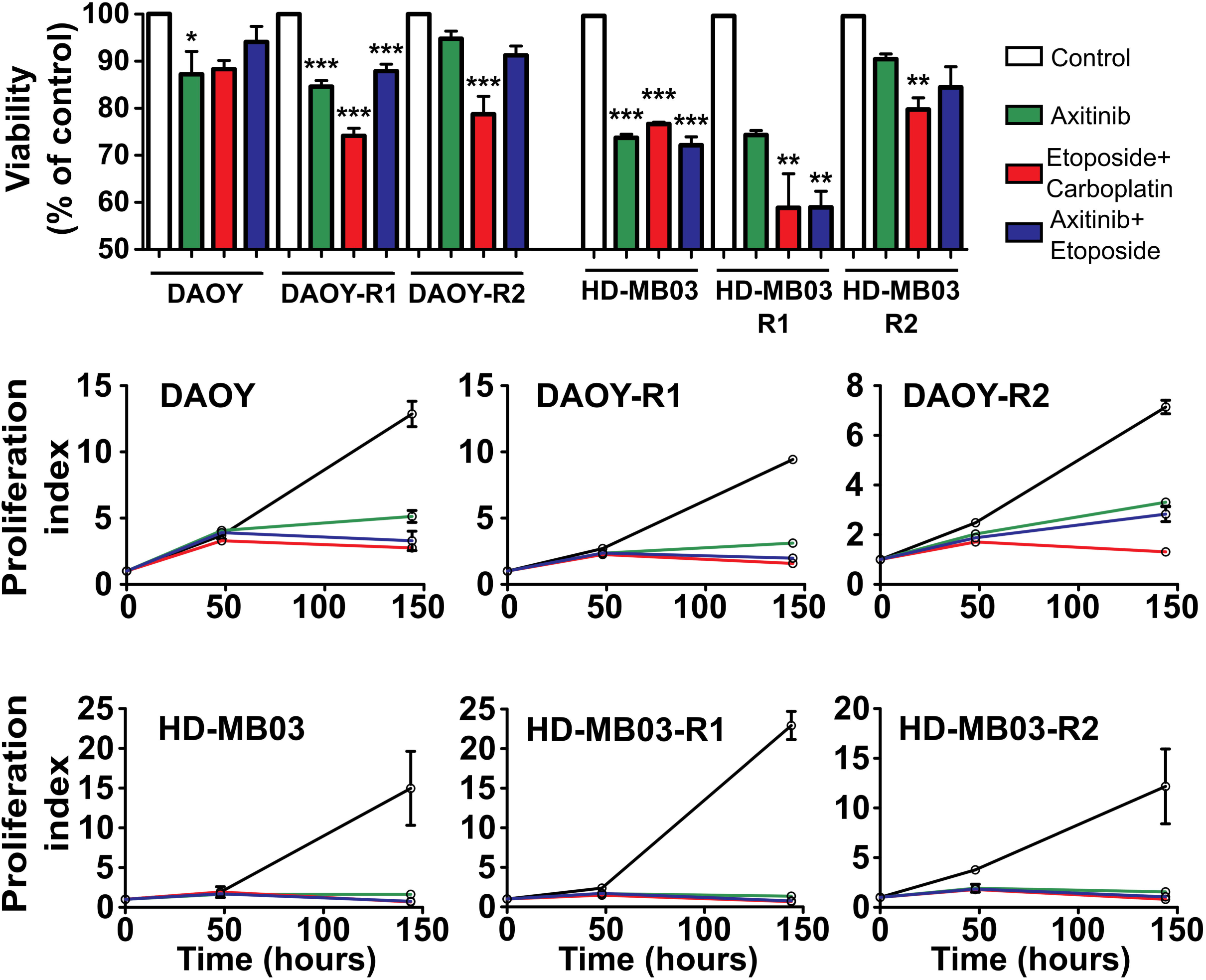
Axitinib and axitinib/etoposide combination reduce the viability and the proliferation of radiation-resistant MB cells. (a) Viability of control (DAOY and HD-MB03) and two independent radiation-resistant (DAOY-R1 and R2 and HD-MB03-R1 and R2) cell populations treated for 48 hours with axitinib (green), an etoposide/carboplatin combination (red) or an axitinib/etoposide combination (blue). All the compounds were dissolved in DMSO, the amount of which was adjusted to be the same in every condition. Control conditions (white) were also treated with the same amount of DMSO. Histograms represent the mean, error bars represent the SEM and each independent data point is represented (white circles); ***: p<0.001, **: p<0.01, *: p<0.05, one-way Anova test, results are not statistically significant unless otherwise stated). (b) Proliferation index of control (DAOY and HD-MB03) and two independent radiation-resistant (DAOY-R1 and R2 and HD-MB03-R1 and R2) cell populations treated for 48 hours with axitinib (green), etoposide/carboplatin combination (red) or axitinib/etoposide combination (blue) (***: p<0.01, one-way Anova test comparing controls to the other experimental conditions).

### Axitinib and axitinib/etoposide combination decrease tumor growth

We performed a tumor xenograft experiment to address the efficiency of axitinib on MB. Importantly, kinases targeted by the axitinib are expressed in MB tumors from patients (Supplementary figure S2). We used cell lines that express high levels of VEGF to generate tumors (Supplementary figure S3). Matrigel plugs containing HD-MB03 (Group 3) or DAOY (Shh group) cells expressing a luciferase reporter gene were injected subcutaneously into nude mice. Tumor engraftment was monitored thanks to luciferase activity. When two consecutive increases in luciferase activity were measured, the tumors were considered to be engrafted, the animals were randomized and the treatments started (Supplementary Fig. S4). Treatments consisted in either high dose axitinib or high dose etoposide (respectively 50mg/kg and 30mg/kg administered orally thrice a week). Moreover, to test the possibility of compensating etoposide dose reduction with axitinib, we treated the animals with a combination of a low dose of axitinib and etoposide (respectively 25mg/kg and 15mg/kg administered three times a week by oral gavage). HD-MB03 control tumors reached 1000 mm3 within 8 to 38 days (Fig.5a). Axitinib and axitinib/etoposide groups however reached the same size after 32 to 63 days and the etoposide group after 21 to 60 days (Fig. 5a). As tumor growth was biphasic with a “lag-time” followed by rapid growth, we applied a linear regression method to analyze the growth rates in the latter phase. We show that tumor growth rates were reduced in mice treated by either axitinib alone or in combination with etoposide when compared to vehicle or etoposide treatments (Fig. 5b and Supplementary Fig. S5).

**Figure 5:**
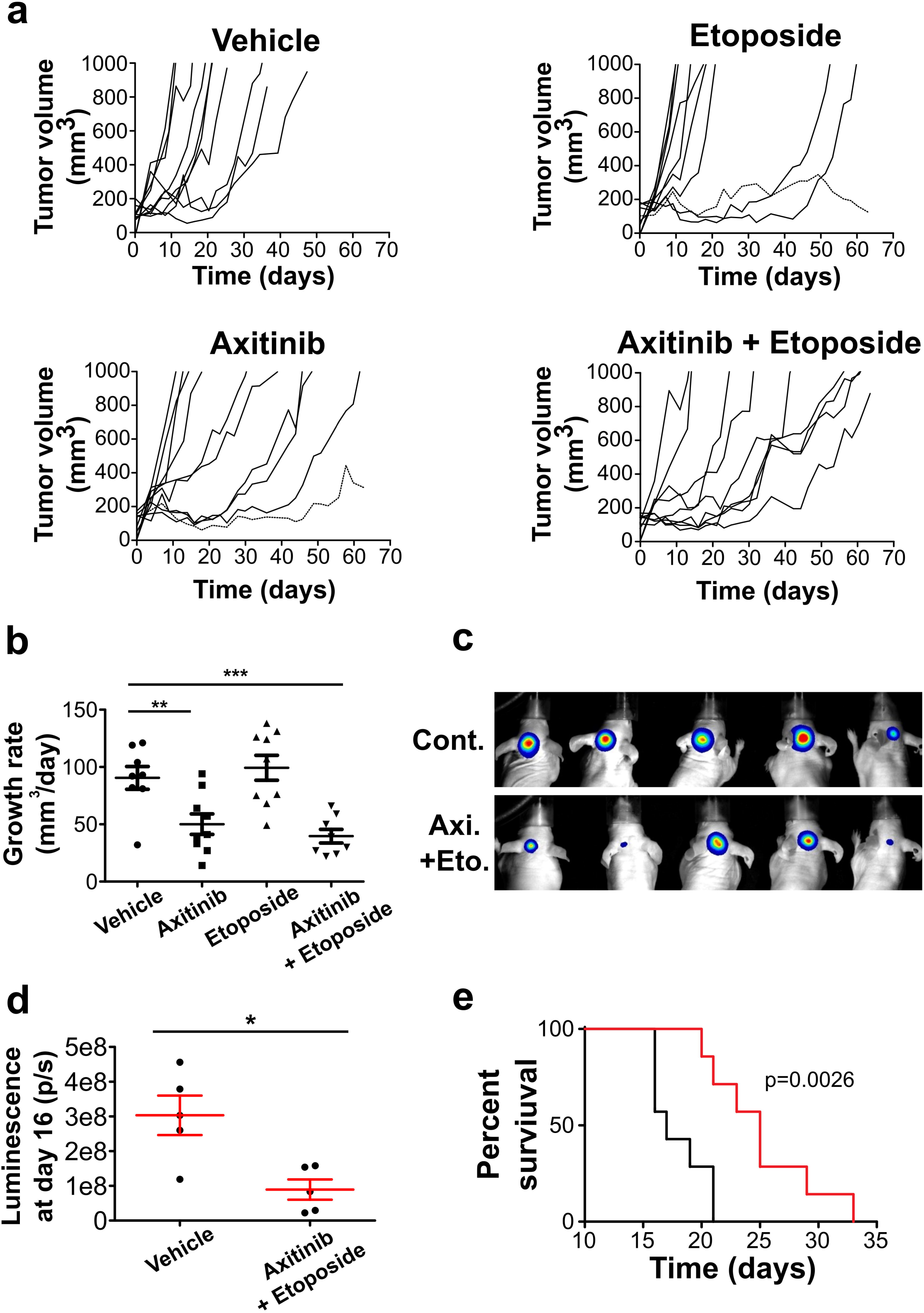
Axitinib and axitinib/etoposide combination reduce tumor growth more efficiently than etoposide. (a) Individual growth curves of subcutaneous HD-MB03 (group 3) tumor xenografts treated with axitinib, etoposide or a combination of axitinib and etoposide (outliers are represented by doted lines). (b) Growth rate of the individual subcutaneous tumors estimated by linear regression (horizontal lines and errors bars represent the mean +/- SEM; ***: p<0.001, **: p<0.01, one-way ANOVA test, results are statistically non-significant unless otherwise stated). (c) Luminescence image of mice after 16 days of orthotopical tumor growth (5 representative mice are presented for each treatment). (d) Quantification of the luminescence of the tumors presented in panel (c) (horizontal lines and errors bars represent the mean +/- SEM; *: p<0.05, Mann-Whitney test). (e) Survival curves of mice orthopically implanted with HD-MB03 in the cerebellum. Day 0 corresponds to the beginning of the treatment (black: control treatment; red: axitinib/etoposide combined treatment, p value of a Log-rank test is indicated).

We also performed a sub-cutaneous xenograft experiment with the Shh group DAOY cells (Supplementary Fig. S6). The lag time preceding tumor growth was more homogenous than in the previous experiment, approximatively ranging from 35 to 45 days (Supplementary Fig. S6a). Mathematical modelling of tumor growth rate suggested that axitinib did not impair tumor growth while etoposide and axitinib/etoposide treatments decreased tumor growth rates (Supplementary Fig. S6b). However, despite no difference in DAOY tumor volume, luminescence measurements indicate that the quantity of tumoral cells was lower after axitinib/etoposide treatment compared to the other treatments (Supplementary Fig. S6c and d). This suggests that the axitinib/etoposide treatment was in fact more efficient than the other treatments against DAOY tumors.

We next performed an orthotopic xenograft experiment to test the efficiency of this combination treatment. HD-MB03 spheroids were implanted in the mouse cerebellum and tumor engraftment was monitored with luciferase activity as in the previous experiment. Treatments consisted in a combination of a low dose of axitinib and etoposide (respectively 25mg/kg and 15mg/kg administered three times a week by oral gavage). The combination treatment strongly reduced tumor growth (Fig. 5c and d) and significantly increased the survival of the animals (Fig. 5e). Taken together, our results suggest that axitinib alone or a combination of lower doses of etoposide and axitinib are efficient *in vivo* on MBs belonging to group 3 and Shh group.

### Axitinib decreases tumor cell proliferation and induces tumor fibrosis

Histological analysis of the subcutaneous tumors revealed that the treatments strongly decreased the number of proliferative Ki67-positive cells both in HD-MB03 and DAOY tumors (Fig. 6a and b and Fig. S7a and b). Axitinib alone or in combination with etoposide induced necrosis in the HD-MB03 tumors (Fig. 6a and c). In these tumors, axitinib or etoposid did not impact the number of endothelial cells (CD31 positive) and/or pericytes/cancer associated fibroblasts (αSMA positive) when compared to controls (Fig. 6a and d). Instead, the axitinib/etoposide combination decreased the number of αSMA and/or CD31positive cells and increased the number of double-stained αSMA/CD31 structures representative of functional vessels covered with pericytes. Masson’s trichrome staining of DAOY tumors showed that every treatment was associated with increased fibrosis (Fig. S7c and d). These results suggest that axitinib mediates its anti-tumor effect via different mechanisms depending on the MB genetic subgroup.

**Figure 6:**
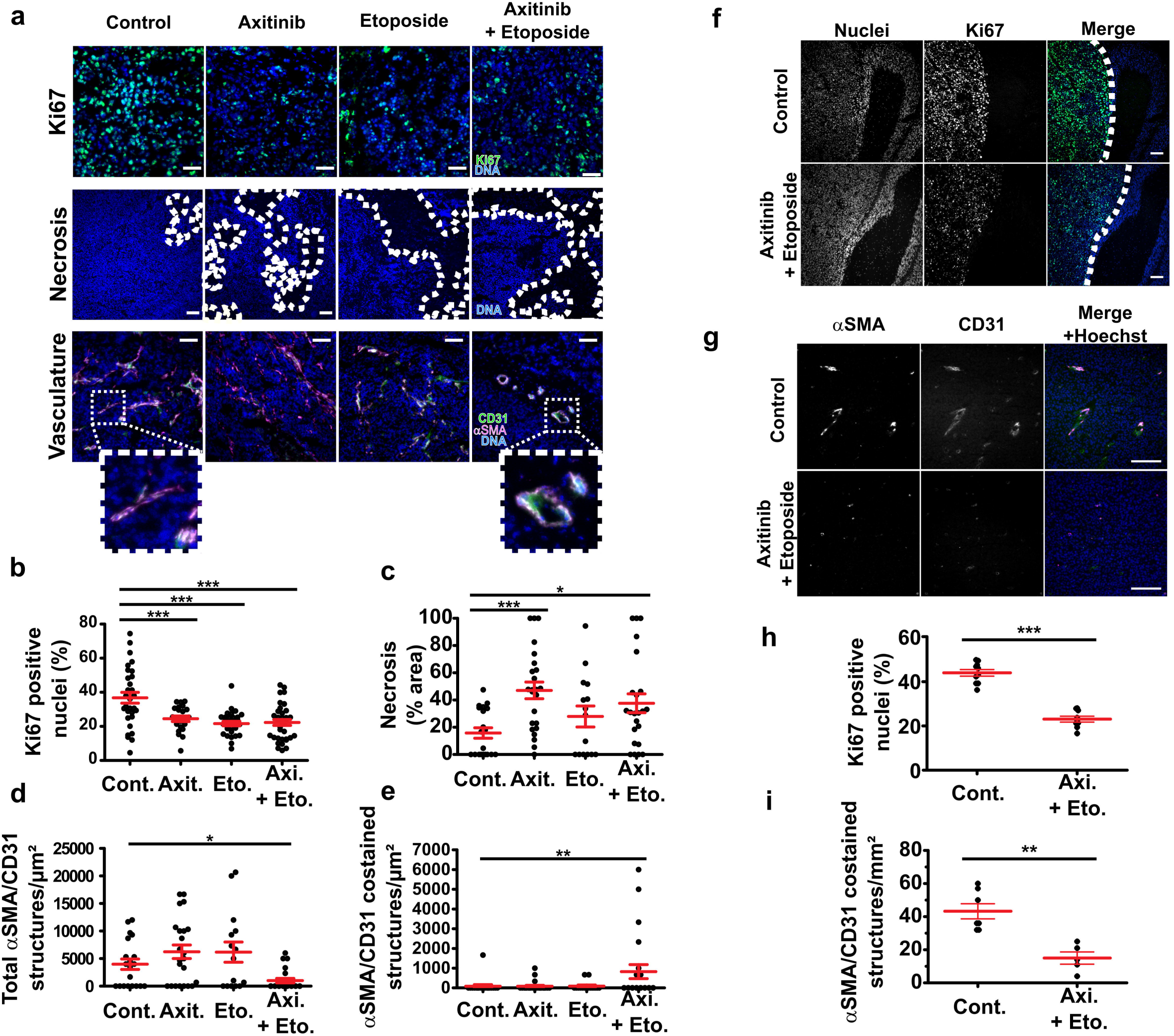
Axitinib, etoposide and axitinib/etoposide treatments reduce cell proliferation and induce tumoral vascularization and necrosis in HDMB tumors. (a-e) Histological analysis of HD-MB03 subcutaneous xenografts. (a) Images of sections of tumors treated with the indicated compounds: proliferative cells revealed by Ki67 immunofluorescent staining (green) and Hoechst33342 nuclear DNA counterstaining (blue); necrosis revealed by DNA counterstaining (blue) (dotted lines delineate necrotic areas); vasculature revealed by CD31 and αSMA immunofluorescent staining (green and magenta respectively) and Hoechst33342 nuclear DNA counterstaining (blue) (images are representative of at least four independent tumors, scale bars: 50µm). (b) Dot plot representing the quantification of the proportion of Ki67 positive nuclei in the indicated experimental conditions. (c) Dot plot representing the quantification of the proportion of necrotic area in the indicated experimental conditions, (d) the number of CD31 and/or αSMA structures per µm^2^ and (e) the number of functional blood vessels per µm^2^ (bars represent the mean +/- SEM; ***: p<0.001, **: p<0.01; *: p<0.05, one-way ANOVA test). (f-i) Histological analysis of HD-MB03 orthotopic xenografts. (f) Images of sections of tumors treated with the indicated compounds. Proliferative cells are revealed by Ki67 immunofluorescent staining (green) and Hoechst33342 nuclear DNA counterstaining (blue) (dotted lines indicate the border between normal and tumoral tissue, scale bars: 200µm). (g) Images of sections of tumors treated with the indicated compounds. The vasculature is revealed by CD31 and αSMA immunofluorescent staining (green and magenta respectively) and Hoechst33342 nuclear DNA counterstaining (blue) (dotted lines delineate the border between normal and tumoral tissues). Images are representative of at least four independent tumors, scale bars: 50µm. (h) Dot plot representing the quantification of the proportion of Ki67 positive nuclei and (i) the number of functional blood vessels per mm^2^ in the indicated experimental conditions (bars represent the mean +/- SEM; ***: p<0.001, **: p<0.01, Mann-Whitney test).

Histological analysis of the HD-MB03 orthotopic tumors showed that the axitinib/etoposide combination significantly decreased the proportion of proliferative Ki67 positive cells in the tumors (Fig. 6f and g). This result suggests that axitinib/etoposide treatment directly impact the proliferation of cancer cells and that this treatment can efficiently cross the blood-brain barrier (BBB).

### Axitinib is not toxic for mammal neonates and can cross the blood-brain barrier

Two important questions regarding the potential use of a new compound to treat pediatric brain tumors are the toxicity of the compound towards developing organisms and its ability to pass the blood-brain barrier (BBB). To address these questions, 20 day old weaning rats were chronically treated daily with 50mg/kg axitinib administered by oral gavage for four weeks. Although no direct correlation can be established between rat and human age, this early stage has been proposed to be comparable with a period around one year old for humans ^23^.

Treatment with axitinib had no effect on the general behavior and growth rate of both female and male animals (Fig. 7a). At the end of the experiment, the animals were sacrificed, the cerebella were collected and their axitinib content was measured. Importantly, we show that axitinib was present in the cerebellar tissue (Fig. 7b). Although a previous report showed that axitinib was flushed out of the brain very efficiently by ABCG2 and ABCB1/2 efflux pumps after an acute exposure ^24, 25^, there is no data regarding the effect of a chronic treatment on axitinib brain accumulation. Thus, in order to get further insight into the mechanism leading to axitinib brain accumulation, we hypothesized that BBB permeabilization may be a result of long-term treatment with axitinib. To test this hypothesis, we used an in vitro model of BBB ^26^. We first determined the toxicity of three doses of axitinib (1, 5 and 25µM) on human brain and umbilical vein endothelial cells (hereafter abbreviated BECs and HUVECs respectively) (Fig. 7c). Axitinib only displayed toxicity on brain endothelial cells after a three day treatment at doses higher than 1µM. We therefore chose the doses of 0.5, 1 and 5µM and a 3 days treatment to test the effect of axitinib on the permeability of our in vitro BBB model (Fig. 7d). We show that axitinib permeabilized the barrier formed by BEC or HUVEC cells at non-toxic doses. These results support the idea that axitinib induces permeabilization of the BBB, allowing it to accumulate in the brain.

**Figure 7:**
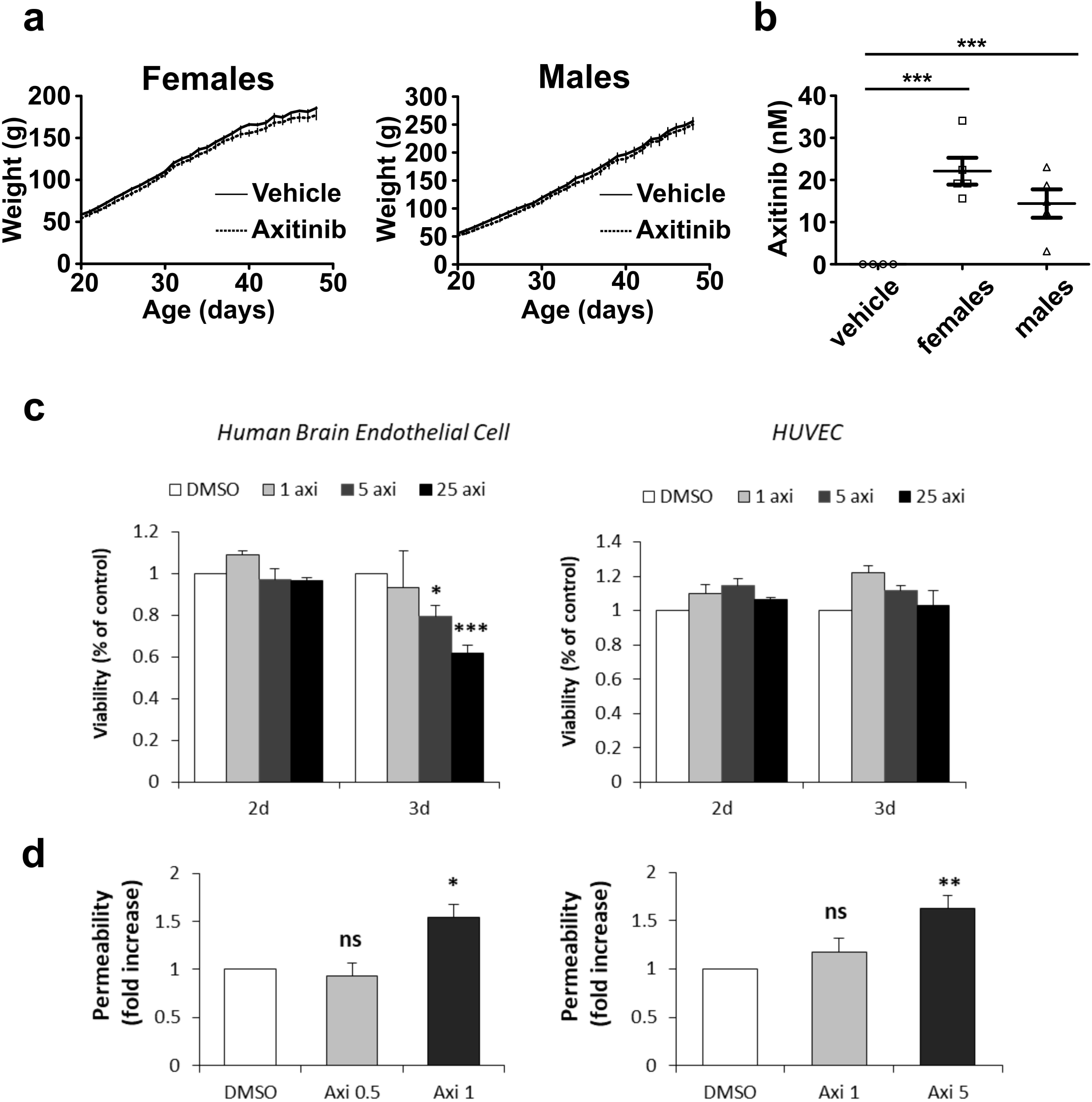
Axitinib shows no major chronic toxicity and can pass the blood-brain-barrier of juvenile mammals. (a) Growth curves of female and male 20 days old Wistar rats treated with axitinib (50mg/kg/day) for 28 days (data points represent mean weigh +/- SEM, n=5 rats per group). (b) Dot plots representing the measurements of axitinib concentrations in rat cerebella after 28 days of treatment (50mg/kg/day Axitinib or vehicle alone, n=5 mice per group, bars represent the mean +/- SEM). (c) MTT assay analysis of axitinib toxicity towards human brain endothelial (BEC) and human umbilical vein endothelial cells (HUVEC). (d) Relative permeability of human brain endothelial cells and HUVEC in an *in vitro* endothelium permeability assay. (*: p<0.05, **: p<0.01, ***: p<0.001, one-way ANOVA test)

### High expression of axitinib targets is linked to poor prognosis in Shh patients

To define the genetic subgroup of patients that may benefit from an axitinib treatment, we correlated the expression of axitinib targets (VEGF receptors 1, 2, 3, cKit and PDGFRA and B) to survival in available data across 763 primary samples of MB, generated by Cavalli and colleagues (Table 3) ^27^. For patients of Group 3, overexpression of cKit and PDGFRA was associated with a shorter survival (p =0.04 and p=0.012, respectively) while overexpression of PDGFRB and VEGFR1 was correlated to a longer survival (p=0.017 and p=0.041, respectively). For patients of Group 4, overexpression of PDGFRB and VEGFR1 is synonymous of shorter survival (p=0.04 and p=5.5E^-3^, respectively) while it is the contrary for VEGFR2 (p=0.043). For Shh patients, rapid death is correlated with overexpression of cKit, PDGFRA, VEGFR1 and VEGFR2 (p=8.7E^-5^, p=6.7E^-5^, p=4.8E^-3^ p= 5.2E^-3^, respectively) while overexpression of PDGFRB and VEGFR3 is linked to a longer survival (p=5E^-8^, p=0.015, respectively). We attributed a relative strength to each parameter, and we gave a “weight” of +2 for a good prognostic marker and a “weight” of -2 for a bad prognostic marker. The worst score was obtained for patients of the Shh subgroup (−5). Therefore, these patients would be good responders to the drug. These in silico results are not in agreement with those obtained with the xenografts experiments (axitinib is more efficient in a model of Group 3 tumors as compared to a model of Shh tumors). These results strongly suggest that the determination of axitinib targets in the primary tumor or at relapse on reference treatment is recommended before administration of the compound.

**Table 3:** Log-rank statistical test associated to the survival curves of the mice bearing subcutaneous HD-MB03 tumor xenografts. Axitinib, Etoposide and Axitinib+Etoposide treatment are associated with a significantly increased survival when compared to the controls.

| Log-rank Test |  |
| --- | --- |
| Control vs Axitinib | ** (p=0.0077) |
| Control vs Etoposide | * (p=0.0483) |
| Control vs Axitinib+Etoposide | ** (p=0.0024) |
| Etoposide vs Axitinib+Etoposide | n.s. (p=0.2139) |
| Axitinib vs Axitinib+Etoposide | n.s. (p=0.4713) |
| Axitinib vs Etoposide | n.s. (p=0.6438) |

## DISCUSSION

In the present study, we show that some anti-angiogenic TKi compounds efficiently kill MB cells belonging to any molecular subgroup. Amongst these compounds, we show that axitinib displays a pleiotropic effect as it is efficient against MB cells from any molecular subgroup and is capable of permeabilizing (in vitro) and crossing (in vivo) the BBB. We further show that axitinib was the compound displaying the lowest toxicity towards non-cancer cells, suggesting it may induce lower toxicity for pediatric patients. Consistent with these results, we show that high dose daily treatments of juvenile rats did not lead to acute toxicity. Finally, we also show that axitinib alone or in combination with reduced dose etoposide is efficient against MB in vivo, both in subcutaneous and cerebral tumors xenografts. As mentioned above, a phase I clinical study has recently established the pharmacokinetics and the maximum tolerated and recommended dose of axitinib in a pediatric context ^28^. This study also pinpoints preliminary evidence of axitinib efficacy in children, although no patient presenting a brain tumor was included in this trial. Treatment with axitinib alone or combined with the PI3K inhibitor GDC-0941 has also showed efficacy on several in vitro MB models ^22, 29^. Taken together with the current work, these studies pave the way towards the use of axitinib to treat pediatric patients with solid tumors of the central nervous system.

We attempted to find which of the described axitinib targets were inhibited upon exposure (data not shown). However, we were unable to identify a mechanism responsible for its toxicity towards MB cells. Indeed, axitinib impairs a wide variety of kinase and non-kinase targets ^29, 30^. Thus, an explanation for our failure in identifying the mechanism responsible for axitinib toxicity towards MB cells probably lies in the fact that this effect is due to combined impairment of several targets at the same time. It would therefore be very difficult to identify single targets of axitinib whose activity is necessary for the survival of MB cells. At the same time, this particular property of targeting many different proteins may also explain the efficacy of axitinib on MB cells and tumors belonging to different molecular subgroups. Moreover, complete molecular profiling of every case of MB everywhere in the world is definitively not a realistic option. Therefore, the use of treatments effective towards any form of MB and not specific to a given molecular subgroup still needs to be seriously considered.

Bringing compounds through the BBB to treat cerebral tumors is extremely challenging. Indeed, the BBB is formed by the walls of the brain capillaries and strictly controls which of the blood flow components can enter the brain compartment ^31, 32^. Endothelial cells of the BBB form a highly impermeable cell layer and express high amounts of efflux pumps that are in charge of exporting molecules out of the brain. Axitinib was showed to be actively removed from the cerebral compartment by the action of the ABCB1 efflux pump system in mice treated with a single dose of the compound ^24, 33^. Nonetheless, axitinib has also proven efficiency against orthotopically implanted mouse glioblastoma ^34^, supporting the idea that it can access brain tumors. Importantly, the formation and maintenance of the BBB was showed to be under the control of the Wnt/β-catenin pathway ^35-37^. Axitinib was recently described to efficiently inhibit the activity of this pathway ^30^. In the present study we also show evidence that this compound permeabilizes an in vitro BBB model and that it can be found in the brain of chronically treated rats. Thus it is tempting to speculate that axitinib permeabilizes the BBB through inactivation of the Wnt/β-catenin pathway. The present study describes the growth inhibition of cerebral xenografts by an axitinib/etoposide combination treatment. This effect could be due to an increased accessibility of the treatments due to the permeabilization of the BBB by the axitinib.

In conclusion, the demonstrated usability of axitinib in a pediatric context together with its predicted low toxicity and ability to target heterogeneous tumors in the brain compartment make it a promising compound to add in the therapeutic arsenal against pediatric brain tumors.

## MATERIAL AND METHODS

### Cell lines

DAOY (ATCC, HTB-186) cells were maintained in MEMα supplemented with 7.5% fetal calf serum (FCS) (Dominique Dutscher SAS), 0.25% Glutamax, 1% NEAA and 0.1% sodium pyruvate (Thermo Fisher Scientific Inc.). D283-Med (ATCC, HTB-185) and D341-Med (ATCC, HTB-187) were cultured in MEM (Thermo Fisher Scientific Inc.) supplemented respectively with 15% fetal calf serum (Dominique Dutscher SAS). CHLA-01-Med (ATCC, CRL-3021) were cultured in DMEM/F12 (Thermo Fisher Scientific Inc.) supplemented with 2% B-27 supplement (Fisher Scientific), 20ng/ml EGF and 20ng/ml basiFGF (Sigma-Aldrich). HD-MB03 (DMSZ, ACC 740) and maintained in RPMI medium (Thermo Fisher Scientific Inc.) supplemented with 7.5% FCS. D458 Med (Cellosaurus, CVCL_1161) and maintained in Improved MEM (Thermo Fisher Scientific Inc.) with 7.5% FCS and 0.25% Glutamax. Human dermal fibroblasts (HDF) (Sigma-Aldrich 106-05N) were maintained in fibroblast growth medium (Sigma-Aldrich). C8-D1A cells (ATCC CRL-2541) were maintained in DMEM (Thermo Fisher Scientific Inc.) supplemented with 7.5% SVF. Immortalised human cerebral microvascular endothelial cells (hCMEC/D3) and Human Umbilical Vein Endothelial Cell (HUVEC) were maintained in Endothelial Basal Medium-2 (EBM-2, Lonza), containing 5% foetalbovine serum (FBS Serum Gold, PAA Laboratories), 1% penicillin/streptomycin (P/S), HEPES, and chemically defined lipid concentrate (Invitrogen), hydrocortisone (1.4 mM), acid ascorbic (5 mg/ml) and basic fibroblast growth factor (1 ng/ml; Sigma) as described by Weksler et al. ^38^. DAOY, HD-MB03 and D458 cell lines were obtained from Dr. Celio Pouponnot’s lab (Institut Curie, Paris, France). D283-Med, D341-Med, CHLA-01-Med and C8-D1A cells were purchased from ATCC. HDF cells were purchased from Sigma-Aldricht. HUVEC cells were purchased from Lonza. The absence of mycoplasma was verified on a bimonthly basis using the PlasmoTest kit (Invivogen, cat. code rep-pt1).

### Lentiviral infections

Lentiviral particles were prepared according to standard protocol as previously described ^39^. Briefly, lentiviral vectors pLenti-CMV-V5-Luc (Addgene plasmid 21474) or pLV-mCherry (Addgene plasmid 36084) were co-transfected with lentivirus packaging vectors psPAX2 (Addgene plasmid 12260) and pMD2.G (Addgene plasmid 12259) into HEK293T cells through PEI transfection. DAOY and HD-MB03 cells (1 and 5 million respectively) were seeded in 100mm diameter culture dishes and incubated with viral supernatant (1:10 V:V) at 37°C overnight. Medium was then replaced with fresh medium. pLV-mCherry stably infected cells were selected for 7 days with 1µg/mL puromycin (InVivoGen).

### MTT, proliferation assays cell viability and cumulative population doubling assays

HDF (3 000 cells), C8-D1A (5 000 cells), D458 (5 000 cells), DAOY (2 000 cells) HD-MB03 (10 000 cells) and CHLA-01-Med (10 000 cells) were seeded in 96 well plates (Corning Inc.) in 100 µl medium per well. A range from 50nM to 150µM of every drug was tested. The EC50 was determined using the non-linear fit function of the Prism 5 software (Graphpad Software Inc.). Alternatively, hCMEC/D3 ^38^ or HUVEC cells were plated at 5000 cells/well and incubated 24h later for 2 and 3 days with 1, 5 or 25µM of Axitinib. The effect of the anti-angiogenic compounds were measured using the 3-[4,5-dimethylthiazol- 2yl]-diphenyltetrazolium bromide (MTT) colorimetric assay (Sigma, Lyon, France) according to the manufacturer’s instructions. When applicable, EC50’s were calculated using the non-linear regression method of Prism 5 software (Graphpad Software Inc.).

Cell viability of cultures treated for 48 hours with 5µM axitinib (Selleck Chemicals S1005), Cabozantinib (Selleck Chemicals S1119), Sunitinib (Selleck Chemicals S7781) or a combination of etoposide (Selleck Chemicals S1225) and carboplatin (Selleck Chemicals S1215) at 1µM and 1.6µM respectively was measured by an propidium iodine incorporation technique using a ADAM-sHIT automated cell counter (NanoEnTek) according to the manufacturer’s protocol.

DAOY, D458, D283, C8-D1A, and HDF (30 000 cells per well) and HD-MB03 (50 000 cells per well) were seeded in 24 wells plates for proliferation assays. Cells were counted the next day (time 0) and after 48 and 120 hours. The proliferation index was determined by dividing the number of cells at a given time by the initial number of cells. Population doublings were calculated as previously described ^40^.

### Analysis of viability and proliferation of radiation-resistant cells

Irradiation-resistant cells were kindly provided by SM (unpublished data). Wild type or two independent cell populations of irradiation-resistant DAOY (DAOY-R1 and DAOY-R2, 30 000 cells per well for proliferation assays and 100 000 cells per well for flow cytometry assays) and HD-MB03 cells (HD-MB03-R1 and HD-MB03-R2, 50 000 cells per well for proliferation assays and 300 000 cells per well for flow cytometry assays) were seeded in 24-wells plates and 6-well plates for proliferation and flow cytometry assays respectively. Cells were treated the next day with 5 µM axitinib (Selleck Chemicals S1005), a combination of etoposide (Selleck Chemicals S1225) and carboplatin (Selleck Chemicals S1215) at 1 µM and 1.6 µM respectively, or a combination of 2,5 µM axitinib and 0,5µM etoposide. For proliferation assays, cells were counted at time 0 and after 48 and 144 hours and the proliferation index were calculated by dividing the number of cells at a given time by the number of cells at time 0. For flow cytometry assays, cells were trypsinized after 48 hours and labeled with 50µg/ml propidium iodide (thermofisher-P3566). Cells labeled were quantified using a Melody FACS (BD Biosciences). Analysis of the FACS data was conducted with Flowjo software (Tree Star, Inc).

### Coculture and FACS experiments

C8-D1A cells were stained with the green fluorescent probe neuro-DiO (Biotium) according to the manufacturer’s protocol. C8-D1A-neuroDio cells (150 000) and HD-MB03-mCherry or DAOY-mCherry cells (50 000) were plated in in 12 wells plates. Cells were exposed to axitinib (5µm) or Eto/Carbo (1µM/1.6µM) treatment during 3 days, the medium was washed and cells were left to recover for 3 days without treatment. Fluorescence images of the cells were taken with a DMI400 (Leica Microsystems) inverted microscope equipped with a 40x objective and a Zyla 5.5 camera (Andor Technologies). Cells were then trypsinized and the number of both cell types was quantified using a Melody FACS (BD Biosciences) with a 488nm laser beam. Analysis of the FACS data was conducted with Flowjo software (Tree Star, Inc).

### Spheroid assays

2 000 cells were seeded in ultra-low adhesion 96 well plates (Corning Inc.). After 4 days, they were transferred in DMEM-7.5% FCS supplemented with 5uM axitinib or 1µM/1.6µM Eto/Carbo and cultured for 8 days. Pictures were taken with an AMG Evos microscope 40x objective (Thermo Fisher Scientific Inc) and the spheroid areas were measured using ImageJ software (NIH, USA).

### Subcutaneous xenografts

For subcutaneous xenograft experiments, tumor cells expressing the luciferase (350 000 HDMB-Luc cells and 1.10^6^ DAOY-Luc cells) were resuspended in 200µL of 5µg/mL matrigel (Corning Inc.) and injected subcutaneously in the flank of 5-week-old Rj:NMRI-Foxn1 nude (nu/nu) female mice (Janvier Labs). 100µL of saline solution (NaCl 0.9M) containing 30 mg/mL D-Luciferin (PerkinElmer) was injected intraperitoneally in the animals and the bioluminescence was quantified using the In Vivo Imaging System (Perkin Elmer) according to the manufacturer’s instructions. Tumor volume (V = L × l^2^ × 0.52) was determined with a caliper. 50 mg/kg axitinib, 30 mg/kg Etoposide (Selleck Chemicals) and a mix of 25 mg/kg axitinib and 15 mg/kg Etoposide resuspended in 200µL of an aqueous solution 0.5% carboxymethylcellulose, 0.4% Tween80 were administered by oral gavage three times a week. Mice were sacrificed when the tumor reached 1000 mm^3^. Tumors that did not reach the size of 500 mm^3^ were considered to be outliers and were not considered during the analysis of the experiments. No statistical methods have been used to predetermine sample size of the experiments. Animal facility availability and cost were used to determine sample sizes. Two strictly independent cohorts of 5 mice per group were used for the tumor growth experiment with HD-MB03 cells and one cohort of five mice per group was used for the experiment with DAOY cells. These experiments were carried out in strict accordance with the recommendations in the Guide for the Care and Use of Laboratory Animals. Our experiments were approved by the ‘‘Direction de l’Action Sanitaire et Sociale” of the Principality of Monaco and the ethic committee of the Centre Scientifique de Monaco.

### Intracranial tumour xenografts

HD-MB03-Luc spheroids were stereotactically implanted into the brains of randomly chosen 9-weeks-old Rj:NMRI-Foxn1 nude (nu/nu) female mice (Janvier Labs). Briefly, MB spheroids (3 spheroids of 2500 cells per mouse) were implanted into the left cerebellar hemisphere (2mm posterior, 1,5mm left of the lambada point and 2,5mm deep) using a Hamilton syringe fitted with a needle (Hamilton, Bonaduz, Switzerland) and following the procedure already described^41^. Mice were treated with 25 mg/kg axitinib and 15 mg/kg Etoposide resuspended in 200µL of an aqueous solution 0.5% carboxymethylcellulose, 0.4% Tween80 administered by oral gavage three times a week. Mouse survival was based on the presence of neuropathological features including gait defects and loss of weight. A minimum of 6 mice per group was chosen to yield enough statistical power (P[=[0.05). These experiments were carried out in strict accordance with the recommendations in the Guide for the Care and Use of Laboratory Animals. Our experiments were approved by the ‘‘Direction de l’Action Sanitaire et Sociale” of the Principality of Monaco and the ethic committee of the Centre Scientifique de Monaco.

### Tumor growth analysis

The growth rate of the tumors generated with HD-MB03 cells was determined by a linear regression method applied to the growth curves between the sizes of 200 mm3 up to the endpoint using Prism 5 (Graphpad Software Inc.). The growth of the tumors generated with DAOY cells could not be fitted to a linear curve and was therefore modeled according to a lognormal model (refer to supplementary methods for details).

### Histopathology and automated image analysis

Tumor samples were recovered from the animals and embedded in OCT compound according to the manufacturer’s protocole (Fisher Healthcare). 5µm thin sections were then prepared with a cryostat (Leica Microsystems). Incubation was carried out with the following antibodies diluted at 1:1000 in TBS supplement with 1% horse serum and 1% BSA for 20 minutes at room temperature: anti-Ki67 (ab16667, Abcam), anti-CD31 (BD Pharmagen #550274) or anti-αSMA (Sigma #A2547). The preparations were then washed with TBS-0.025% Triton, incubated with an anti-rabbit Alexa488 and an anti-mouse Alex-555-coupled secondary antibody (Cell Signaling Technology), washed with TBS-0.025% Triton and the nuclei were counterstained with Hoechst33342 (Thermo Fisher Scientific). Fluorescence images of the cells were taken with a DMI400 (Leica Microsystems) inverted microscope equipped with a 40x objective (Leica Microsystems) and a Zyla 5.5 camera (Andor Technologies) driven by Micromanager software^42^. At least one cross-section of non-overlapping images of each tumor was acquired. All the images were then quantified with CellProfiler 3.0 software ^43^. The pipeline used for this analysis is available from https://github.com/TTteam-CSM/image_analysis.

### Juvenile rats toxicity experiment

Toxicity experiments were carried out on 20 days old RjHan:WI – Wistar rats (10 males and 10 females) treated with 50mg/kg axitinib resuspended in 100µL of a 0.5% carboxymethylcellulose, 0.4% Tween80 aqueous solution on a daily basis by oral gavage. All experiments were serviced by Janvier Labs Company (Saint Berthevin, France) and conducted at the Janvier Labs facility in strict accordance with the recommendations in the Guide for the Care and Use of Laboratory Animals. The experiments were approved by the French “Comité National Institutionnel d’Ethique pour l’Animal de Laboratoire”.

### BBB permeability assay

This assay was conducted as previously described ^26^. Briefly, hCMEC/D3 or HUVEC cells were plated at 250.000 cells/inserts and incubated 24h later for 3 days with 0.5 or 1µM Axitinib. Permeability was measured by the passage of FITC-Dextran 40kDa through the inserts. Fluorescence intensity measures were normalized to DMSO control.

### Statistical analyses

All experiments were carried out in technical triplicates and separately repeated at least twice (the detailed number of replicates are available in the figure legends). One-way ANOVA test was used to measure significance between several experiment groups. Student’s t-tests were used to measure significance between control and an experiment group when indicated in the figure legends. Data were analyzed with Prism 5 software (Graphpad Software Inc.). The tests were performed with a nominal significant level of 0.05. *p < 0.05, **p < 0.01, and **p < 0.001. Results are showed as mean ± SEM.

### Data availability statement

The datasets generated and analysed during the current study are available from the corresponding author on reasonable request

## ACKNOWLEDGMENTS

This work was supported by the Government of the Principalty of Monaco and a grant from Fondation Flavien, Un Nouvel Espoir. MPB was funded by Fondation Flavien, Un Nouvel Espoir. The authors acknowledge Dr. Patrick Gizzi and TechMedIll for the analysis of axitinib content in the rat cerebella. The authors are extremely grateful to all the staff members of Fondation Flavien for their constant efforts and strong support. The authors warmly thank Dr. M. Gettings for editing the manuscript and

## AUTHOR CONTRIBUTIONS

VP and MPB designed and performed the experiments. MPB, GP, VV and VP analysed the data. VP wrote the manuscript and all the co□authors contributed in improving the draft manuscript. SM provided the irradiation-resistant MB cells. JDo. contributed to design the study and to improve the manuscript. MPB, JDu., AV, VV, VP and RG performed and VPB contributed to the modeling and statistical analysis of the tumor xenograft experiments. VV performed the histological experiments. VV and AV provided technical assistance. JG performed the in vitro BBB permeability assay.

## COMPETING INTERESTS STATEMENT

The authors declare no competing interests.

## Notes

### Competing Interest Statement

The authors have declared no competing interest.

