## Supplementary material for "“The anti-angiogenic compound axitinib demonstrates low toxicity and anti-tumoral effects against medulloblastoma”"

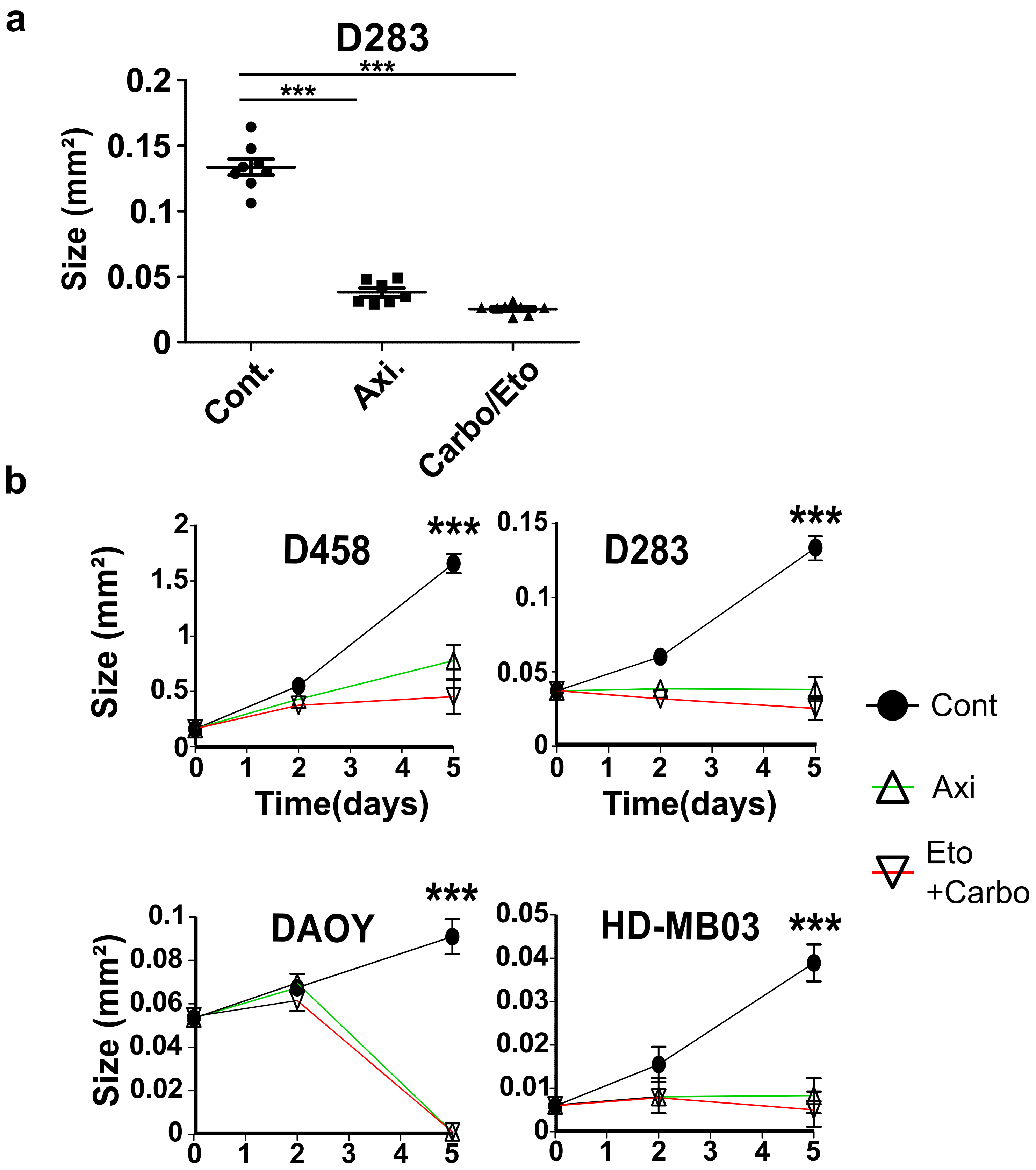


**Supplementary Figure S1: Axitinib and axitinib/etoposide combination reduce the growth of MB spheroids more efficiently than etoposide alone.** (a) Dotplot showing the endpoint size measurements of spheroids generated with the D283 MB cell line and continuously treated with 5μM axitinib (Axi.), a combination of 1μM etoposide and 1.6μM carboplatin (Carbo/Eto). Controls were all treated with a concentration of DMSO corresponding to the one used as vehicle for the drugs. (b) Growth curves of spheroids generated with the indicated cell lines and treated as indicated above (results presented are from at least 3 independent experiments, ***: $p<0.001$, one-way ANOVA test).

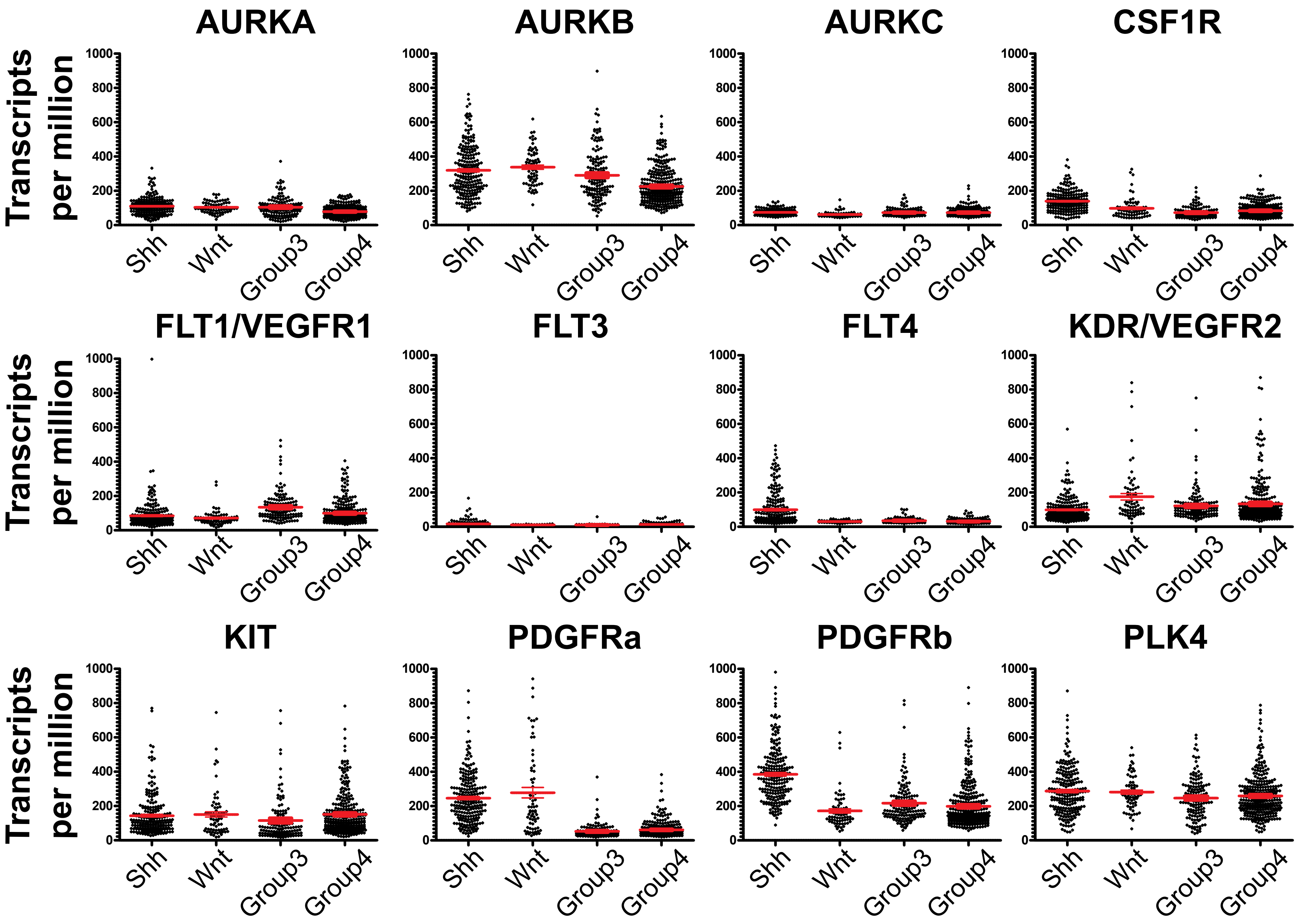


**Supplementary Figure S2: Expression of axitinib kinase targets in MB cases.** The expression level of axitinib described kinase targets across the four subgroups of MB was extracted from the transcriptomic data obtained by Cavalli and colleagues[27].

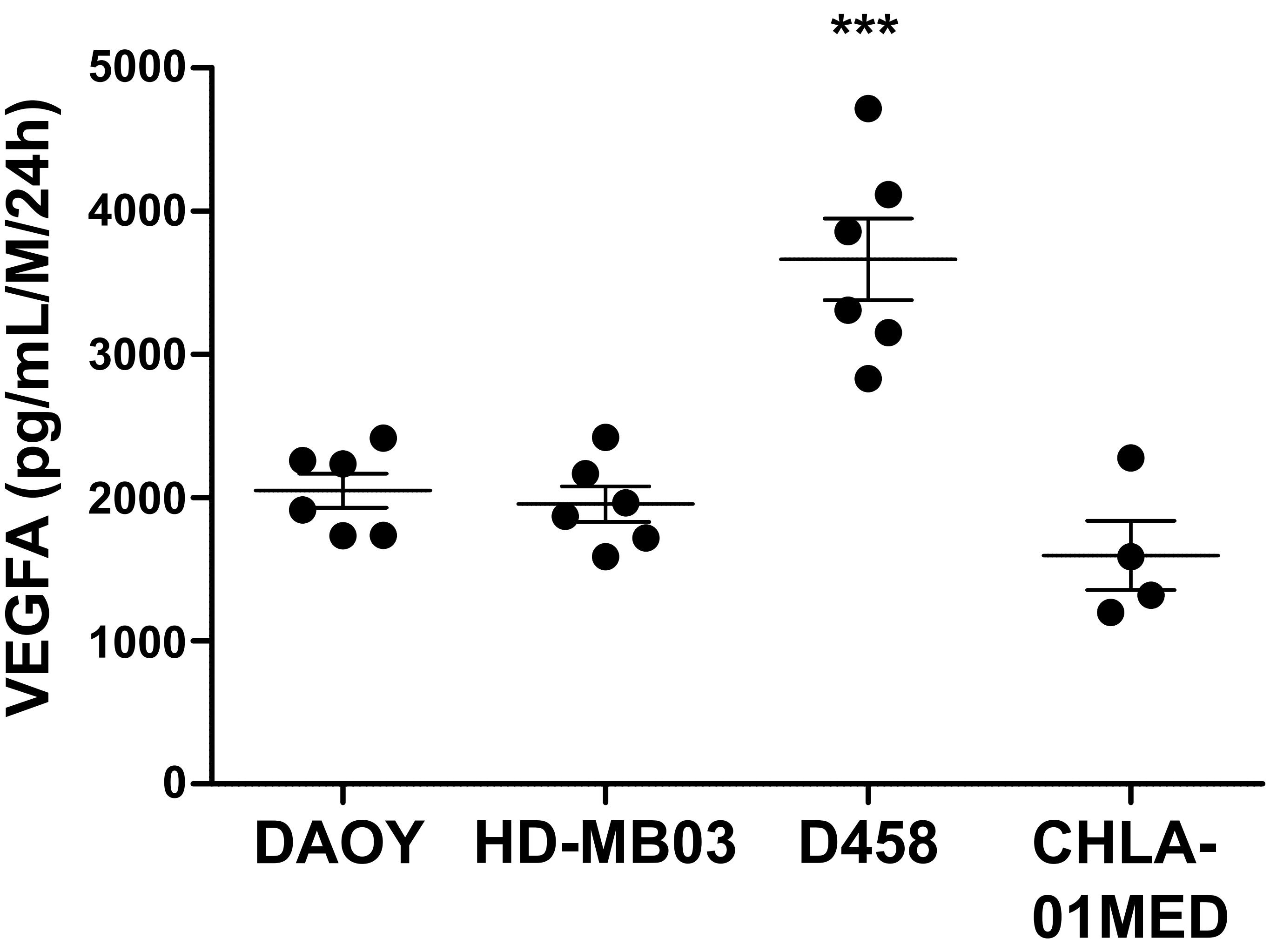


S**upplementary Figure S3: VEGFa expression in MB cell lines.** Dotplot presenting the expression level of VEGFa measured by ELISA in 2D cultures of four MB cell lines. The concentration level of VEGFa is expressed in pg per mL per million cells per 24 hours. (results presented are from at least 3 independent experiments, bars represent the mean +/- SEM, ***: p<0.001, one-way ANOVA test).

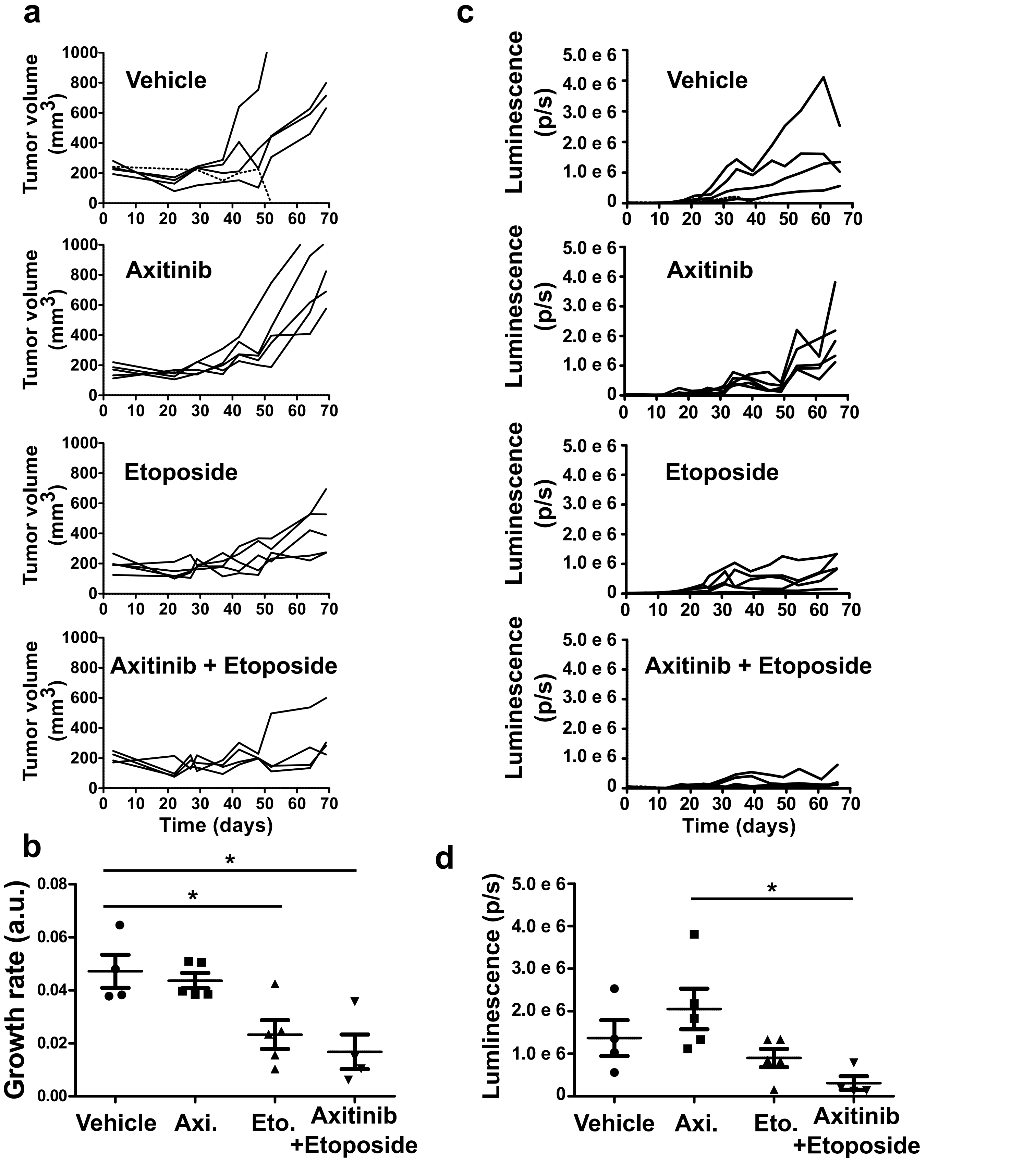


**Supplementary Figure S4: Axitinib/etoposide combination reduces tumor growth more efficiently than etoposide alone.** (A) Individual growth curves of subcutaneous DAOY (Shh group) tumor xenografts treated with Axitinib, Etoposide or a combination of both (outliers are represented by dotted lines). (B) Relative growth rates for each condition were determined by mathematical modelling. The median growth rate and 97.5% confidence interval are represented. (C) Individual curves representing the luminescence of subcutaneous DAOY (Shh group) tumor xenografts treated with Axitinib, Etoposide or a combination of both (outliers are represented by doted lines). (D) Dot plot presenting the endpoint luminescence levels for each treatment (bars represent the mean +/- SEM, *: $p<0.05$, one-way ANOVA test, results are non-statistically significant unless otherwise stated).

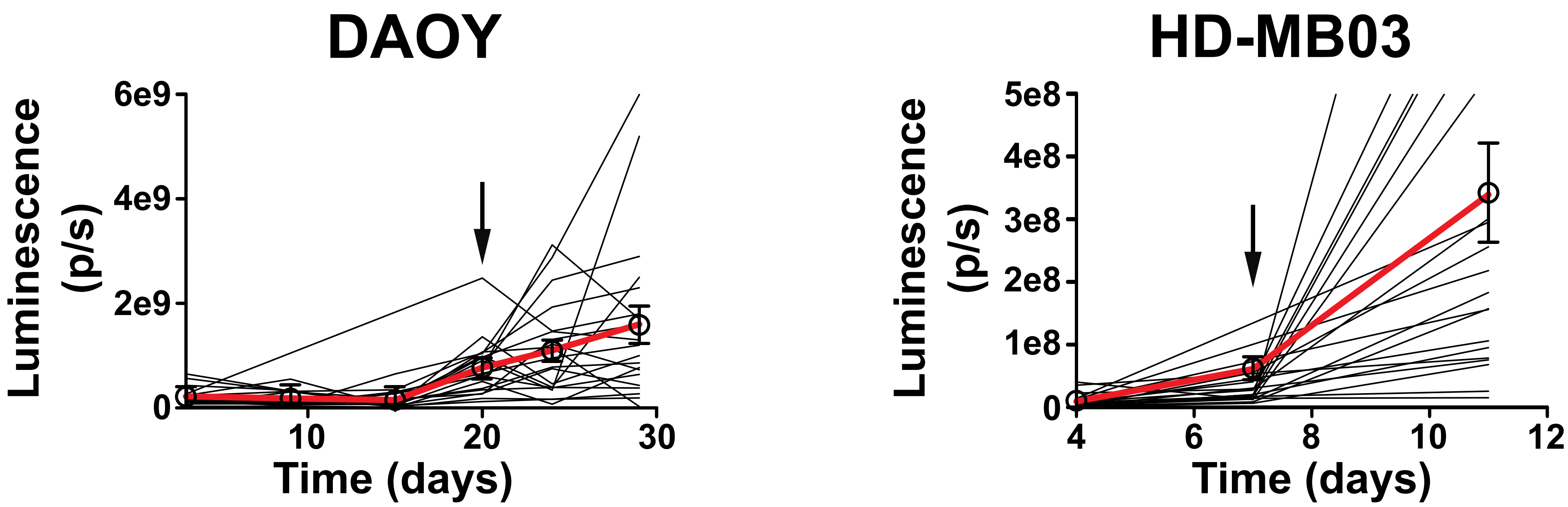


**Supplementary Figure S5: Tumor growth rate evaluation by a linear regression method.** Growth curves were generated for each individual tumor starting from the first measurement over 200mm$^3$ and a linear regression method was used to estimate the average growth rate of each tumor (estimated slopes+/- SEM and for the corresponding fits $R^2$ are presented in the table aside the corresponding growth curves).

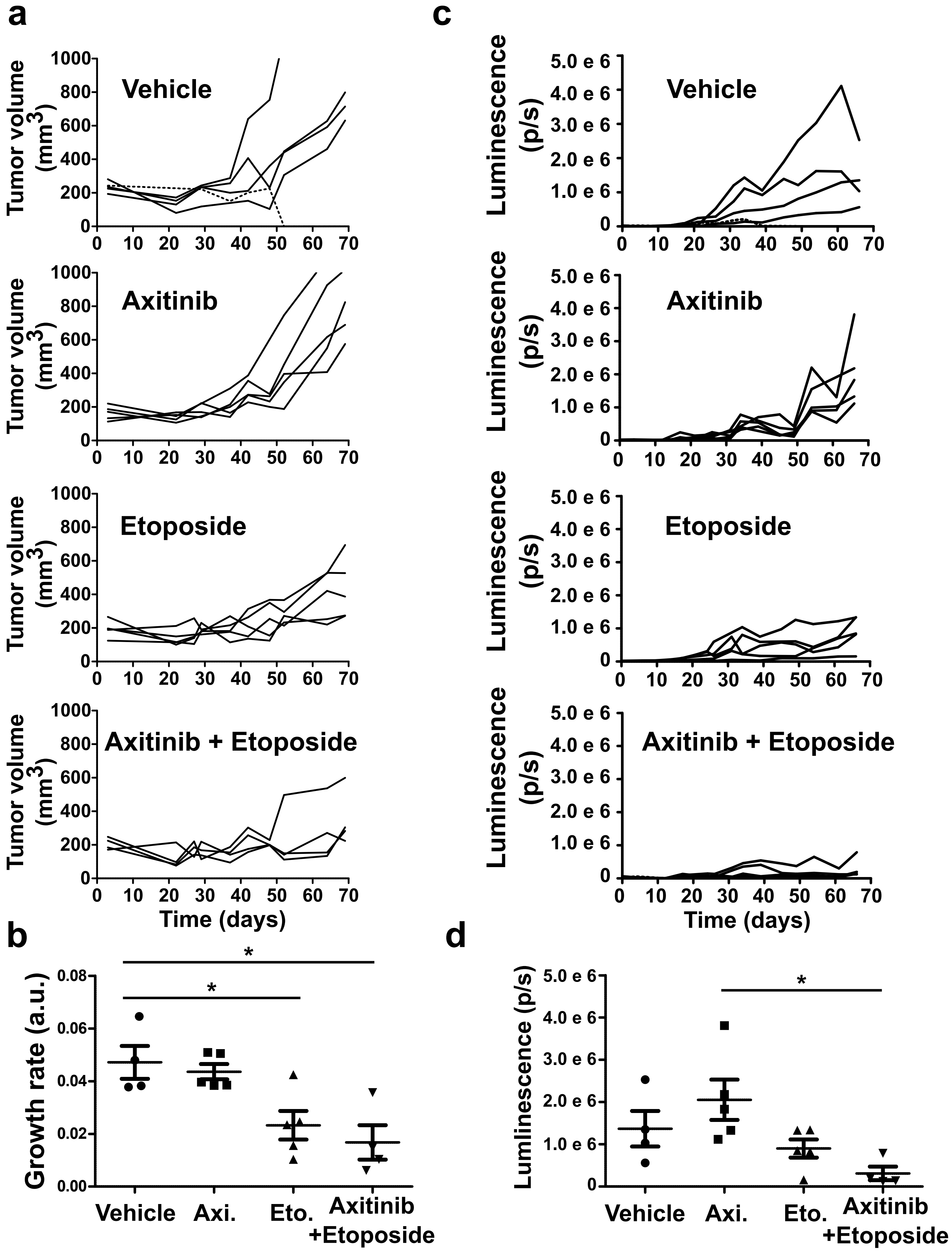


**Supplementary Figure S6: Axitinib/etoposide combination reduces tumor growth more efficiently than etoposide alone.** (a) Individual growth curves of subcutaneous DAOY (Shh group) tumor xenografts treated with Axitinib, Etoposide or a combination of both (outliers are represented by dotted lines). (b) Relative growth rates for each condition were determined by mathematical modelling. The median growth rate and 97.5% confidence interval are represented. (c) Individual curves representing the luminescence of subcutaneous DAOY (Shh group) tumor xenografts treated with Axitinib, Etoposide or a combination of both (outliers are represented by doted lines). (d) Dot plot presenting the endpoint luminescence levels for each treatment (bars represent the mean +/- SEM, *: $p<0.05$, one-way ANOVA test, results are non-statistically significant unless otherwise stated).

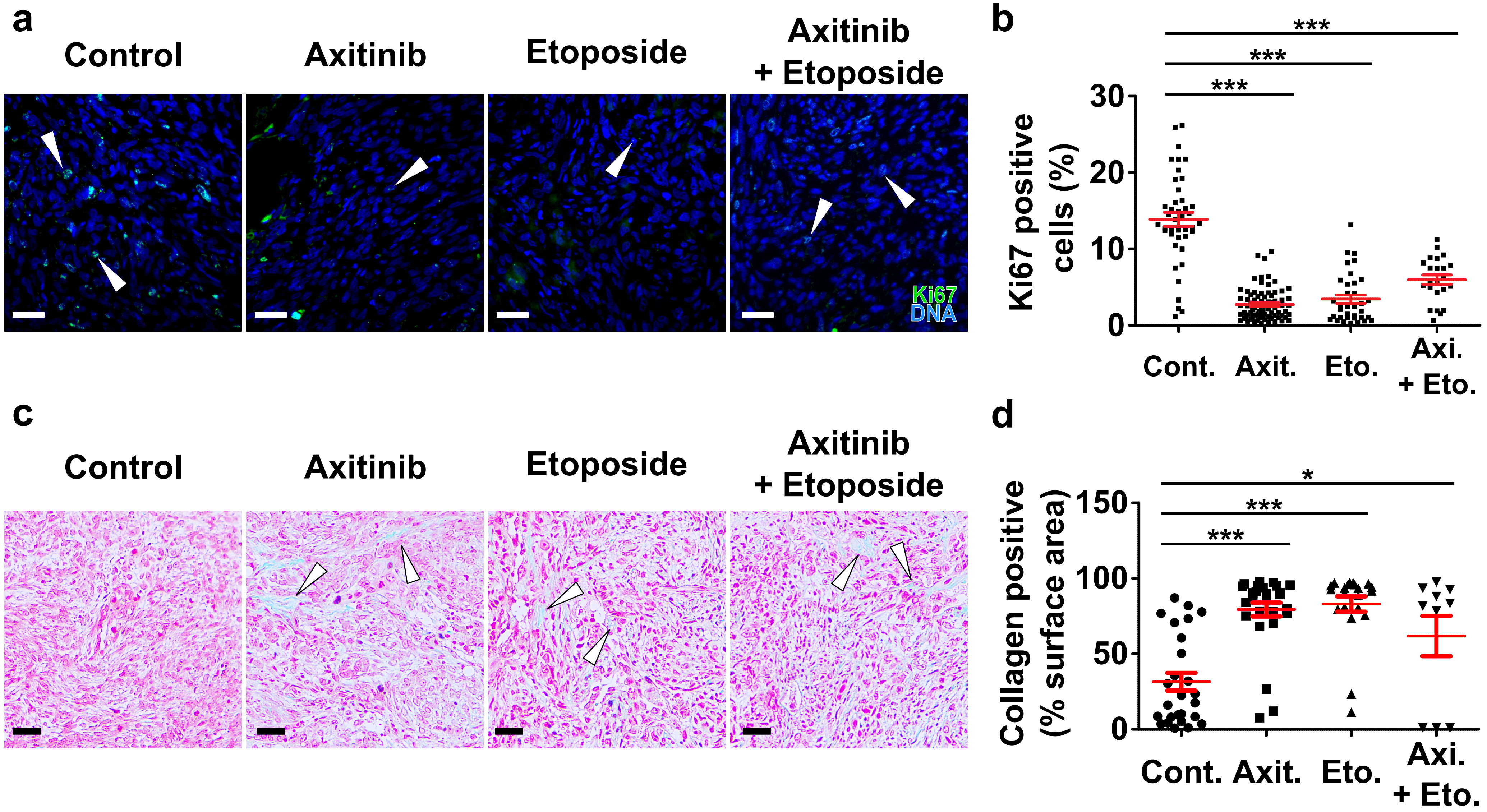


**Supplementary Figure S7: Axitinib, etoposide and axitinib/etoposide treatments reduce cell proliferation and induce tumoral fibrosis in DAOY tumors.** (a) Ki67 immunofluorescent staining (green) and Hoechst33342 nuclear DNA counterstaining (blue) of sections of DAOY tumor xenografts treated with the indicated compounds. White arrowheads indicate Ki67 positive nuclei. (b) Quantification of the proportion of Ki67 positive cells in the indicated experimental conditions. (c) Masson's trichrome staining (nuclei: dark pink; cytoplasm: light pink, collagen: light blue) of DAOY tumor xenografts treated with the indicated compounds. White arrowheads indicate collagen positive areas. (d) Dot plots representing the proportion of the image fields displaying consistent collagen staining (bars represent the mean +/- SEM). (Images are representative of at least four independent tumors; ***: $p<0.001$, *: $p<0.01$, one-way ANOVA tests; scale bars: 100µm).
